# Microbiota-Mediated Competition Between *Drosophila* Species

**DOI:** 10.1101/2020.08.05.238055

**Authors:** Antoine Rombaut, Romain Gallet, Kenza Qitout, Mukherjy Samy, Robin Guilhot, Pauline Ghirardini, Brian P. Lazzaro, Paul G. Becher, Anne Xuéreb, Patricia Gibert, Simon Fellous

## Abstract

Species that share resources often avoid competition with context-dependent behaviors. This is the case for the invasive insect pest *Drosophila suzukii*, whose larval ecological niche overlaps with that of *Drosophila melanogaster* in ripe, but not rotten, fruit. We discovered *D. suzukii* females prevent costly interspecific larval competition by avoiding oviposition on substrates previously visited by *D. melanogaster*. More precisely, *D. melanogaster* association with gut bacteria of the genus *Lactobacillus* triggers *D. suzukii* avoidance. However, *D. suzukii* avoidance behavior is condition-dependent, and *D. suzukii* females that themselves carry *D. melanogaster* bacteria stop avoiding sites visited by *D. melanogaster*. The adaptive significance of avoiding cues from the competitor’s microbiota was revealed by experimentally reproducing in-fruit larval competition: reduced survival of D. suzukii larvae was dependent on the presence of gut bacteria in the competitor. This study unveils a new role for the symbiotic microbiota and plastic behaviors in mediating interspecific competition.

## Introduction

Over the last 10 years, the Asian fly *Drosophila suzukii* (*Ds*) has spread in Europe and the Americas (1) causing major fruit-production losses (2-4). As a consequence, considerable research effort has been devoted to development of strategies to control this species and protect crops. It was observed that co-culturing of *Ds* with *D. melanogaster* (*Dmel*) led to rapid competitive exclusion of *Ds* (5). This phenomenon can be partly explained by the observation that *Ds* females avoid laying eggs in resource sites that already contain *Dmel* eggs (6). The prevention of larval crowding does however not explain this behaviour as *Ds* females did not avoid oviposition on sites with conspecific *Ds* eggs in (6) and in the conditions of our experiments (Fig. S1). The literature on *Ds* however reports both oviposition preference and avoidance of sites with cues from conspecifics, possibliy because of context-dependency of this response (7, 8). We hypothesized that *Dmel* eggs might carry specific cues that deter *Ds* females from depositing eggs. We investigated the mechanisms and variability of *Dmel* repellency on *Ds* oviposition. We determined that the oviposition deterrence is mediated by *Dmel* symbiotic bacteria, and that the repellency is plastic and conditional on the *Ds* carrying a microbiota distinct from that of *Dmel*. We infer that the inhibition of *Ds* oviposition is a microbiota-mediated adaptive response to reduce larval competition between the two species.

## Results and discussion

### *Variable response of* D. suzukii *females* to D. melanogaster *cues*

In an initial experiment, we offered groups of *Ds* females the choice to oviposit either on substrates previously exposed for 24h to *Dmel* females or on control substrates (Fig. 1). We followed *Ds* egg-laying preferences over four days with the oviposition substrates replaced daily. During the first two days, *Ds* females laid more than 75% of their eggs on sites that had not been exposed to *Dmel* (*p* < 0.01; Fig 2a). However, *Ds* females did not avoid substrate contaminated by *Dmel* during the final two days of the assay. Avoidance of conspecific cues as reported by (8) was temporary too, but disappeared much faster, after 4h in choice conditions. The presented experiment showed that *Ds* have a strong preference to oviposit on sites that have not been visited by *Dmel*, but that the avoidance behavior is plastic and depends on female experience or condition.

**Figure 1:**
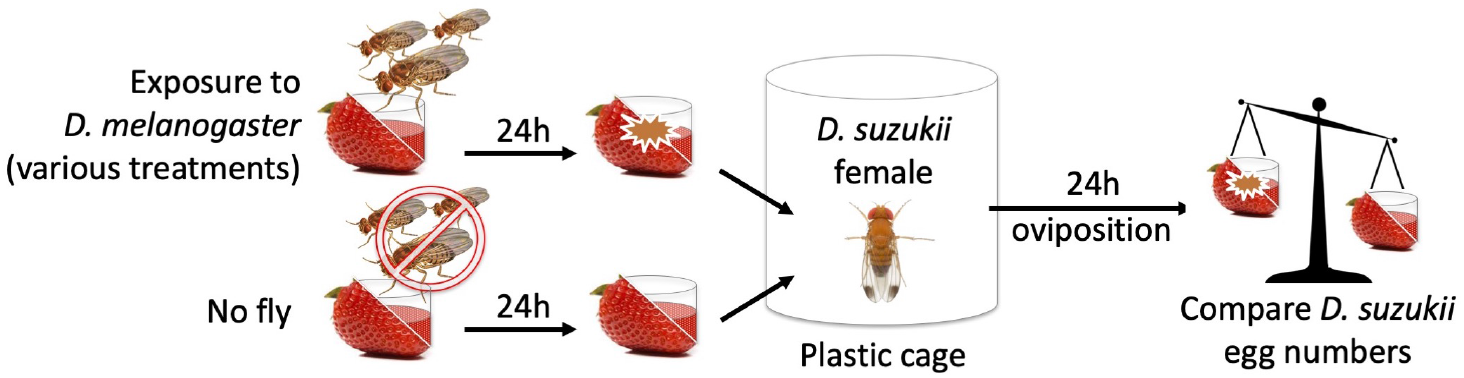
Schematic drawing of the experimental procedure for testing *D. suzukii* oviposition avoidance of egg-laying sites previously exposed to *D. melanogaster*. Details of each experiment, among which 155 origin, sex and numbers of *D. suzukii* and *D. melanogaster* flies, cage size and oviposition substrate, are described in Table 1 of the Materials and Methods.

**Figure 2:**
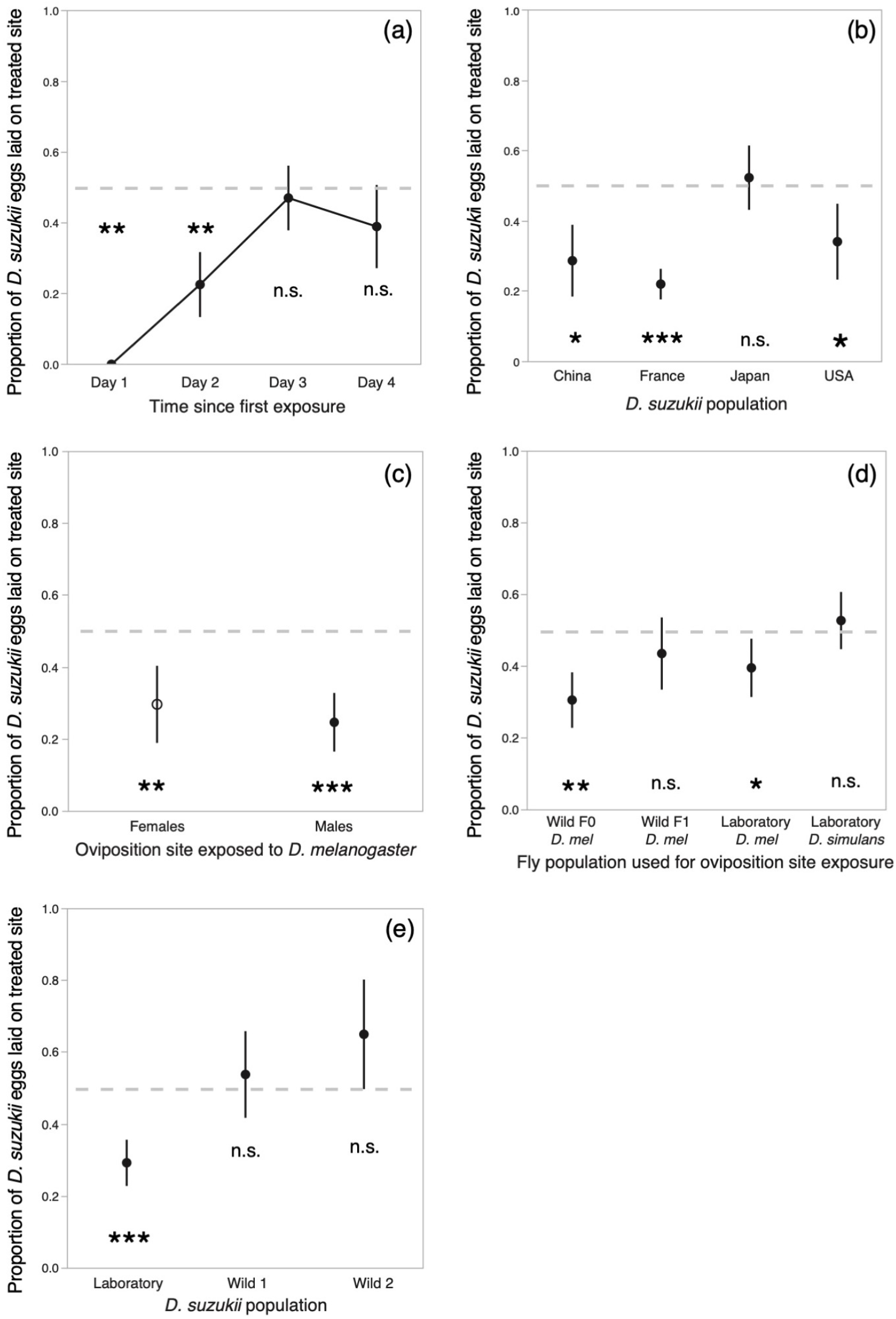
Oviposition avoidance of *D. suzukii* females for egg-laying site previously exposed to *D*. 160 *melanogaster*. Values significantly below 0.5 indicate *D. suzukii* preference for sites unexposed to *D. melanogaster*. Repeated tests of the same females (a) showed plastic avoidance loss. *D. suzukii* populations from different geographical origins (b) exhibited variable avoidance. *D. melanogaster* males, like females, (c) induce repellency. Trap-captured, wild *D. melanogaster* flies (F0 in d) induced repellency, however this property was not induced by laboratory-reared offspring from wild-caught flies (F1 in d) nor by *D. simulans*. Trap-captured, wild *D. suzukii* females (e) did not avoid oviposition on *D. melanogaster* exposed substrates. Symbols indicate means and error-bars standard errors. Significant deviation from equal number of eggs on sites exposed to *D. melanogaster*, or control sites, were produced by one-tailed Wilcoxon signed rank tests; * for p< 0.05; ** for p< 0.01; *** for p< 0.001.

In order to determine how universal, the *Ds* avoidance behavior is, we tested *Ds* females from different laboratory populations founded with insects captured in France (our reference population used throughout this study), the USA, China and Japan (see methods). Because in these behavioral investigations individual females were the essential unit of replication, and so as to consider potential inter-individual differences, this and all following experiments were carried out with single females, rather than groups of flies, and over 24h. Females from all populations except the Japanese exhibited significant avoidance of oviposition on substrates that were visited by *Dmel* (Fig. 2b). *Ds* originates from mainland China, invaded Japan at the beginning of the 20^th^ century, and invaded Europe and North-America in the past 10-15 years (9). These results show that avoidance behavior is neither restricted to invasive populations nor to those from the area of origin.

**Table 1:** details of the experiments presented in Figure 2.

| Experiment<br>question and figure<br>presenting the results | Method | Number of *Ds* females in<br>assay; type of container | *Ds* particulars | *Dmel* particulars | Oviposition substrate | Replication and raw statistical results<br>Reported number of replicates excludes the frequent cases<br>where no *Ds* eggs were deposited during the experiment | Additional comments |
| --- | --- | --- | --- | --- | --- | --- | --- |
| - Do *Ds* females<br>maintain avoidance<br>behaviour over time?<br>- Fig. 2a | We repeatedly assayed the behaviour<br>of the same females over 4<br>consecutive days renewing the<br>oviposition substrates daily. | - Initially 10 females, some<br>mortality occurred during<br>the 4 days of the<br>experiment<br>- 30cm diameter cylinders | Standard laboratory<br>population | Standard Oregon R<br>laboratory<br>population | - Jellified grape juice<br>- 5cm petri-dishes | We followed 9 cages throughout the 4 days.<br>**Wilcoxon signed rank tests, one-tailed:**<br>Day 1: $S=21$; $p=0.008$<br>Day 2: $S=20.5$; $p=0.006$<br>Day 3: $S=6$; $p=0.28$<br>Day 4: $S=9$; $p=0.18$ | On days 1 and 3 *Ds* females were given the choice between *Dme*-<br>exposed and control substrates. On day 2 and 4 they were given the<br>choice between *Dmel*- and *Ds*-exposed substrates. |
| - Is avoidance behaviour<br>present in various *Ds*<br>populations?<br>- Fig. 2b | We assayed avoidance behaviour in 4<br>laboratory populations of different<br>regional origin | - 1 female per assay<br>- 9 cm diameter cylinders | Standard laboratory<br>population and 3<br>populations founded<br>from individuals<br>captured in China, the<br>USA and Japan | Standard Oregon R<br>laboratory<br>population | - strawberry puree<br>- 2*2 cm cubic receptacles | **Wilcoxon signed rank tests, one-tailed:**<br>China: $n=14$, $S=32.5$; $p=0.019$<br>France: $n=57$, $S=557$; $p<0.0001$<br>Japan: $n=27$, $S=-3.5$; $p=0.53$<br>USA: $n=16$, $S=42$; $p=0.014$ | Details of the tested *Ds* populations are available in the biological<br>material section |
| - Are male *Dmel*<br>repellent?<br>- Fig. 2c | Oviposition substrates exposed to<br>either males or females *Dmel*. | - 1 female per assay<br>- 9cm diameter cylinders | Standard laboratory<br>population | Standard Oregon R<br>laboratory<br>population | - jellified grape juice<br>- 2*2cm cubic receptacles | **Wilcoxon signed rank tests, one-tailed:**<br>Female *Dmel*: $n=16$, $S=54$; $p=0.0013$<br>Male *Dmel*: $n=21$, $S=83.5$; $p=0.0007$ | When *Ds* oviposition substrates had been exposed to *Dmel* females that<br>oviposited too, eggs from each species could be discriminated thanks to<br>the elongated respiratory tubes that are specific to *Ds*. |
| - Are wild *Dmel* and<br>laboratory *D. simulans*<br>repellent?<br>- Fig. 2d | We captured wild adult *Dmel* and<br>tested their repellency. We also tested<br>the repellency of F1 offspring from<br>wild-*Dmel*. A *D. simulans* population<br>laboratory was included in the assay | - 1 female per assay<br>- 9cm diameter cylinders | Standard laboratory<br>population | Laboratory and wild<br>*Dmel* ; offspring of<br>wild *Dmel* flies (i.e.<br>F1) ; laboratory *D.*<br>*simulans* population. | - jellified grape juice<br>- 2*2cm cubic receptacles | **Wilcoxon signed rank tests, one-tailed:**<br>Wild F0: $n=17$, $S=57.5$; $p=0.0032$<br>Wild F1: $n=16$, $S=6.5$; $p=0.40$<br>Laboratory: $n=23$, $S=62.5$; $p=0.027$<br>*D. simulans*: $n=19$, $S=7$; $p=0.40$ | Details of the tested *Dmel* populations are available in the biological<br>material section |
| - Do wild *Ds* females<br>avoid *Dmel* exposed<br>media?<br>- Fig. 2e | We captured wild adult flies and<br>assayed their behaviour in the<br>laboratory. | - 1 female per assay<br>- 9cm diameter cylinders | Laboratory and wild flies | Wild *Dmel* | - strawberry puree<br>- 2*2cm cubic receptacles | **Wilcoxon signed rank tests, one-tailed:**<br>Laboratory: $n=19$, $S=74$; $p=0.0008$<br>Wild 1: $n=12$, $S=6.5$; $p=0.28$<br>Wild 2: $n=8$, $S=-15$; $p=0.98$ | Wild *Ds* females were trap-captured in two localities near Montpellier,<br>France. Traps usually contained a diversity of fly species, including *Dmel*.<br>*Ds* females were kept in the laboratory for several days before they<br>started laying eggs and could be assayed. |

Our initial experiments demonstrated that *Ds* females actively avoid oviposition on substrates that had been previously visited by *Dmel* females (Fig. 2a and 2b). But these experiments do not distinguish whether the aversion is due to the presence of *Dmel* flies or *Dmel* eggs. To test this, we repeated the repellency assay using substrate conditioned by *Dmel* males. The experiment showed that *Dmel* males induce the same level of oviposition avoidance as *Dmel* females (p<0.001; Fig. 2c). This rules-out *Dmel* eggs or oviposition-associated cues as driving *Ds* oviposition avoidance, and contrasts with Tephritid fruit flies that use host-marking pheromones to limit oviposition and avoid larval crowding (10).

Because our initial experiments were performed using a laboratory population of *Dmel*, we wanted to determine whether *Ds* oviposition avoidance could also be triggered by wild *Dmel* and by the *Dmel* sister species, *D. simulans* (*Dsim*), whose ecology is very close to that of *Dmel* (11). We tested the repellency of wild *Dmel* trap-captured in Southern France, lab-reared F1 offspring of the same wild Southern France *Dmel* population, the Oregon-R lab strain of *Dmel* used for all previous experiments. Similar to the experiments performed with laboratory *Dmel*, substrate conditioned by the wild *Dmel* flies was repellent to *Ds* females (p<0.01, Fig 2d). Surprisingly, however, the F1 offspring of the wild-caught *Dmel*, which had spent one generation in the laboratory, did not induce oviposition avoidance (Fig. 2d). The *Dsim* population we tested also was not repellent. Similarly, exposure of fruit to *Ds* did not elicit *Ds* oviposition avoidance (Fig. S1). Repellency is therefore a feature of wild and laboratory *Dmel* populations that may nonetheless be sensitive to rearing conditions.

Finally, we tested whether wild *Ds* also avoid substrates that have been visited by *Dmel*. We trapped wild *Ds* adults from the Montpellier region, Southern France, using classical vinegar traps modified to prevent the drowning of captured flies (see methods). These traps attracted various species of Drosophilid flies, including both *Ds* and *Dmel*. To our surprise, wild *Ds* females did not exhibit avoidance behavior to *Dmel*-exposed substrates (Fig. 2e). We can envision three alternative explanations for this: (1) avoidance behavior is a laboratory artefact; (2) uncontrolled fly age or pre-capture history affects female selectivity; or (3) exposure to other Drosophilid flies, including *Dmel*, during the time spent in traps eliminates the avoidance behavior, similar to the third and fourth days our first experiment (Fig. 2a).

Our results show that *Ds* oviposition avoidance of sites with *Dmel* cues varies among populations and with individual experience or physiological condition. This contrasts with the sustained and hard-wired oviposition avoidance that *Dmel* females display in response to geosmin (12), a molecule produced by microorganisms responsible for late-stage fruit rot that are detrimental to *Dmel* larvae. Unveiling the nature of the *Dmel* cues perceived by *Ds* females may shed light on how *Ds* females lose their aversive response.

### Bacterial symbionts of *Dmel* are involved in repellency and *Ds* avoidance loss

Our observation that exposure to males or females of *Dmel* was sufficient to reduce *Ds* oviposition (Fig. 2c) but that *Ds* avoidance behavior was lost after 2 days of exposure to *Dmel* (Fig. 2a) led us to hypothesize that the repellent agent was something shed by all adult *Dm*. To test whether hypothesized agent was volatile or stationary, we conducted an additional experiment testing whether repellency was restricted to substrates directly contacted by *Dmel*-exposed or whether adjacent substrate also became repellent to *Ds*. We did not observe *Ds* avoidance to substrates neighboring *Dmel*-exposed medium (Fig. S2), so we concluded that repellent agent could not diffuse through air. A logical alternative was that *Dmel* might condition the substrate with bacteria they shed, and that the bacteria were aversive to *Ds*. Drosophilids possess the sensory and neuronal circuitry to perceive specific bacteria and compounds produced by them, and the presence of microbiota on substrate has previously been shown to affect behaviors in *Dmel* such as adult foraging preferences (13, 14). Furthermore, the effect of substrate microbes on the behavior of *Dmel* depends on their endogenous microbiota (13, 14). We thus hypothesized that microbial symbionts of *Dmel* excreted on the substrate could perhaps be perceived by *Ds* females, and that oviposition avoidance, or its lack, could be a function of the symbiont community carried by *Ds*.

As a first test of this hypotheses, we experimentally removed the microbiota from *Dmel* and tested whether these axenic flies remained repellent to *Ds*. Because we suspected the *Ds* microbiota might also influence oviposition avoidance, we performed this test with both axenic and conventionally reared *Ds* females (Table 2). Axenic *Dmel* flies did not elicit oviposition avoidance in *Ds*, and both axenic and conventional *Ds* were significantly repelled by conventionally reared *Dmel* (p < 0.01, Fig. 3a). Thus, we conclude that some component of the *Dmel* microbiota is directly or indirectly required to repel *Ds* but that *Ds* does not require microbiota to perceive repellent cues.

**Table 2:** Table 1: details of the experiments presented in Figure 3.

| Experiment<br>question and figure presenting the results | Method | Number of *Ds*<br>females in assay;<br>type of container | *Ds* particulars | *Dmel* particulars | Oviposition substrate | Replication and raw statistical results<br>Reported number of replicates excludes the frequent cases where no *Ds* eggs were deposited during the experiment | Additional comments |
| --- | --- | --- | --- | --- | --- | --- | --- |
| - Is *Dmel* repellence exerted at a distance?<br>- Fig. S2 | We tested whether medium surrounding medium exposed to *Dmel* received less *Ds* eggs than medium surrounding pristine medium. | - 1 female per assay<br>- 20cm cubic netting cages | Standard laboratory population | Standard Oregon R laboratory population | - strawberry puree<br>- 2*2cm cubic receptacles and 12cm square petri dishes | **Wilcoxon signed rank tests, one-tailed:**<br>Exposed area: n= 29, S= 101; p= 0.013<br>Peripheral area: n= 29, S= -14; p= 0.61 | Details of the protocol are described in the supplementary materials |
| - Do axenic *Ds* and *Dmel* flies maintain avoidance and repellence?<br>- Fig. 3a | We produced axenic *Ds* and *Dmel*, which we compared to conventional flies. They were tested in a full-factorial set-up. | - 1 female per assay<br>- 9cm diameter cylinders | Standard laboratory population;<br>conventional and axenic | Standard Oregon R laboratory population;<br>conventional and axenic | - strawberry puree<br>- 2*2cm cubic receptacles | **Wilcoxon signed rank tests, one-tailed:**<br>Conventional *Dmel* and conventional *Ds*: n= 27, S= 133; p= 0.0001<br>Conventional *Dmel* and axenic *Ds*: n= 17, S= 60.5; p= 0.002<br>Axenic *Dmel* and conventional *Ds*: n= 16, S= 19.5; p= 0.11<br>Axenic *Dmel* and axenic *Ds*: n= 16, S= 8.5; p= 0.39 | Details for the production of axenic flies are described in the appropriate section |
| - Test of bacterial candidates possibly involved in *Dmel* repellence<br>- Fig. 4b | We mono-associated axenic *Dmel* flies (i.e. made gnotobiotic flies) with two of their most important bacterial gut reported in the literature, *Lactobacillus brevis* and *Acetobacter pomorum*. We also tested *Dmel* associated with *Escherichia coli*. | - 1 female per assay<br>- 9cm diameter cylinders | Standard laboratory population | Axenic and mono-associated Oregon R flies | - Pieces of bleached strawberries maintained in agar jelly<br>- 2*2cm cubic receptacles | **Wilcoxon signed rank tests, one-tailed:**<br>*L. brevis*: n= 14, S= 40.5; p= 0.0043<br>*A. pomorum*: n= 13, S= 22; p= 0.0745<br>*E. coli*: n=11, S= 10.5; p= 0.18<br>Axenics: n= 17, S= 31; p= 0.071 | Details for the production of axenic and mono-associated flies are described in the appropriate section |
| - Do *Ds* females that bear repellence bacteria still avoid oviposition in sites exposed to *Dmel* associated with the same bacteria?<br>- Fig. 4c | We compared the behaviour of axenic *Ds* females to that of conspecific associated with the same bacterium as in *Dmel* | - 1 female per assay<br>- 9cm diameter cylinders | Axenic and mono-associated with *Lactobacillus brevis* | Oregon R flies mono-associated to *Lactobacillus brevis* | - strawberry puree<br>- 2*2cm cubic receptacles | **Wilcoxon signed rank tests, one-tailed:**<br>Axenics: n= 29, S= 147; p= 0.0003<br>Associated to *L. brevis*: n= 28, S= 54; p= 0.11 | Details for the production of axenic and mono-associated flies are described in the appropriate section |
| - Are bacteria alone sufficient to elicit *Ds* avoidance?<br>- Fig. 4d | We deposited cells of the bacterium *L. brevis* on oviposition medium, let it rest overnight as for *Dmel* exposure, and tested whether it elicited *Ds* avoidance. This assay was repeated with high and low cell numbers, the latter corresponding to the number of cells retrieved on medium surface after *Dmel* exposure in the standard conditions of our experiments. | - 1 female per assay<br>- 9cm diameter cylinders | Axenics | mono-associated Oregon R flies (i.e. bacteria in *Dmel*) and pure bacteria from liquid culture | - strawberry puree<br>- 2*2cm cubic receptacles | **HIGH bacterial dose (1,000,000):**<br>**Wilcoxon signed rank tests, one-tailed:**<br>*Dmel* exposure: n= 29, S= 147; p= 0.0003<br>Bacteria only: n= 29, S= 103; p= 0.011<br>**LOW bacterial dose (5,000):**<br>**Wilcoxon signed rank tests, one-tailed:**<br>*Dmel* exposure: n= 22, S= 82.5; p= 0.002<br>Bacteria only: n= 11, S= 2.5; p= 0.58 | Details for the production of axenic, mono-associated flies and purified bacteria are described in the appropriate section |

**Figure 3:**
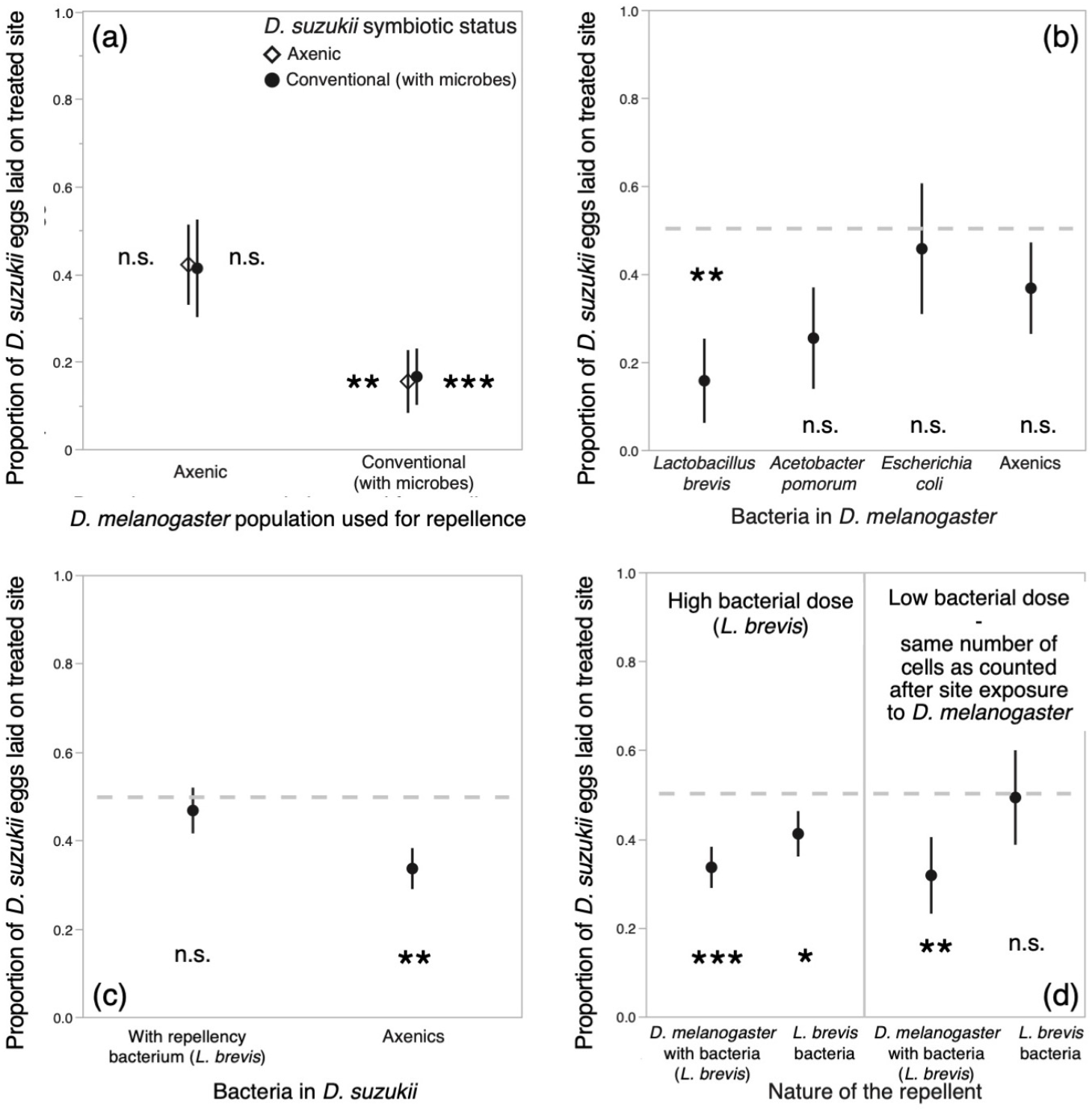
Investigation of the role of extracellular symbionts on *D. melanogaster* repellence and *D. suzukii* oviposition avoidance. Axeny, the removal of extra-cellular microorganisms, (a) had different effects on *D. melanogaster* and *D. suzukii*. Oviposition sites exposed to axenic *D. melanogaster* were not avoided by *D. suzukii*, showing the importance of symbionts in *D. melanogaster* for repellence. By contrast, axenic *D. suzukii* behaved like conventionally reared flies; *D. suzukii* microorganisms were therefore not required for perceiving the repellent. Tests of candidate bacteria in association with *D. melanogaster* (b) revealed the bacterium *Lactobacillus brevis* can restore repellence in formerly axenic flies (note the axenic and *Acetobacter pomorum* treatments were marginally non-significant, p= 0.071 and p= 0.075, respectively). We hypothesized *D. suzukii* avoidance loss was due to their colonisation with *D. melanogaster* symbionts. As expected, *D. suzukii* females experimentally associated with the bacterium *L. brevis* (c) did not avoid oviposition on sites exposed to *L. brevis*-associated *D. melanogaster*. Direct inoculation of medium with *L. brevis* cells (d) in large numbers or at a dose similar to that naturally shed by *D. melanogaster* (i.e. 1,000,00 *vs* 5,000) produced different results. The low, natural dose of deposited bacteria failed to elicit avoidance, suggesting *D. melanogaster* repellence is largely due to the production of unidentified molecules when in symbiosis. Symbols indicate means and error-bars standard errors. Statistical tests produced by Wilcoxon signed rank tests; * for p< 0.05; ** for p< 0.01; *** for p< 0.001.

To identify the specific bacteria responsible for generating repellence in *Dmel*, we inoculated axenic flies with candidate bacteria (i.e. creating gnotobiotic flies). The bacterial microbiota of *Dmel* has been extensively described over the last ten years, showing it largely varies among populations and environmental conditions but almost always includes species of the genera *Lactobacillus* and *Acetobacter* (15-18). We therefore elected to associate *Dmel* flies with a strain of *Lactobacillus brevis*, or with one of *Acetobacter pomorum*, both of which had been isolated from a laboratory population of *Dmel* and are frequently used for microbiota studies (19, 20). In order to test whether any generic bacterium could restore repellency in axenic *Dmel*, we also associated *Dmel* flies with a strain of *Escherichia coli* previously shown as non-pathogenic to flies (21). *Dmel* inoculation with *L. brevis* made *Dmel* repellent to *Ds* (p < 0.01) while association with *A. pomorum* and *E. coli* did not (Fig. 3b). This experiment identifies *L. brevis* as a bacterium able to induce *Dmel* repellency and demonstrates that repellency is not common to all bacteria. It is possible that differences in the microbiota explain the different repellency intensities we observed in the different experiments we conducted. Determining the full range of microorganisms able to render *Dmel* repellent will require further work with wild microorganisms and flies under conditions likely to be experienced in the field.

In our initial experiments, we observed that *Ds* females lose avoidance behaviour after two days of exposure to *Dmel* cues (Fig. 2a). How to explain this change? We hypothesized that the decrease of oviposition avoidance was due to colonisation of *Ds* by the microorganisms deposited by *Dmel* on oviposition sites. In order to test this hypothesis, we mono-associated adult *Ds* for 5 days with the strain of *L. brevis* that elicited strong repellence by *Dmel* (Fig 3b). As expected, *Ds* females associated with *L. brevis* did not avoid oviposition on substrate that had been exposed to *Dmel* adults bearing the same bacterium (Fig 3c). *Ds* females hence avoided sites with cues indicative of presence of *Dmel* unless they carried similar bacteria. This result could also explain why trap-captured wild *Ds* females did not avoid *Dmel* cues (Fig. 2e). In the traps, wild *Ds* were in close contact with other Drosophilids from which they may have acquired microbiota.

To investigate the possibility of transferring our results to application in pest management, we investigated whether bacteria deposited by *Dmel* were sufficient to repel *Ds* oviposition even in absence of *Dmel* individuals, or if *Ds* flies perceive cues produced by the interaction between *Dmel* and its symbionts. A recent study indeed shows *Ds* females respond to bacterial contamination and avoid oviposition in sites inoculated with bacteria-rich wash-water from *Dmel*-exposed media (22). To investigate the effect of *L. brevis* inoculation we carried out two experiments. In the first, we tested the repellence of medium inoculated with 1,000,000 *L. brevis* bacterial cells. In the second, we inoculated the medium with only 5,000 cells, which corresponds to the approximate number of live bacteria sustained on substrates exposed to *Dmel* under our experimental conditions. *Ds* females avoided oviposition on media inoculated with the larger number of bacterial cells (p < 0.05; Fig 3d), although not as strongly as they 240 avoided substrates exposed to *Dmel* flies. *Ds* did not avoid oviposition on substrates inoculated with the lower number of *L. brevis* (Fig. 3d). Together, these results suggest that when *Dmel* adults are associated with bacteria, the interaction produces compounds that are shed and perceived by *Ds* females, but that neither the *Dmel* fly nor her associated bacteria are sufficient for full repellency on their own. A recent study reported that bacteria deposited during oviposition by the oriental fruit-fly, *Bactrocera dorsalis*, induce the host fruit to produce a molecule, b-caryophyllene, that is perceived by female flies and repels them from ovipositing (23). In the case of *Ds* ovipositional avoidance, prospects for crop protection will necessitate identifying the compounds produced by the interaction between *Dmel* and its bacteria and testing them as pure molecules.

### *Ds* larvae suffer from competition with symbiont-associated *Dmel* larvae

Avoidance behavior by *Ds* females could be an adaptation that ensures offspring do not develop in poor quality sites. In order to test whether *Ds* larvae suffer from competition with *Dmel* larvae, we reproduced in-fruit competition between the two species. Surface-sterilized grape berries were pierced with a fine-needle and single *Ds* eggs deposited in each hole, mimicking *Ds* oviposition (Fig. S3a). Holes received axenic or conventional *Dmel* eggs, and, using a full-factorial design, we deposited either 1 or 5 *Dmel* eggs per *Ds* egg. The number of eggs of each species were based on infestation intensities observed in field-collected fruit from which both species had emerged (24). *Ds* developmental success (i.e. proportion of eggs that reached adulthood) was impaired by competition with *Dmel* larvae that were associated with their microbiota, but not with axenic *Dmel* larvae (Fig. 4). In the wild, *Dmel* eggs are never axenic, so the normal outcome of larval competition should therefore be poor *Ds* development. These results support our hypothesis that *Ds* oviposition behavior prevents costly larval competition with *Dmel*. Our results however contrast with those of Bing (25) who observed *Ds* larvae suffer from the presence of *Dmel* bacteria such as *L. brevis*. Here, the presence of *Dmel* bacteria in absence of *Dmel* larvae did not reduce *Ds* larval survival (Fig. 4, left). We inoculated the fruit through exposure to *Dmel* males so it is possible that the bacteria did not reach *Ds* larvae high numbers, especially since *Lactobacillus* is predominantly anaerobic. However, our data show unambiguously that the combination of *Dmel* larvae and their microbiota is detrimental to *Ds* development. Whether *Ds* larvae suffered directly from bacterial presence, from direct interactions with microbiota-associated *Dmel* larvae, or from metabolic byproducts of the *Dmel*-microbiota association is unknown. Each of these mechanisms is plausible, and gut-bacteria effects on *Drosophila* larvae and antagonistic interactions among *Drosophila* larvae are environment dependent (e.g. 20, 25, 26, 27, 28).

**Figure 4:**
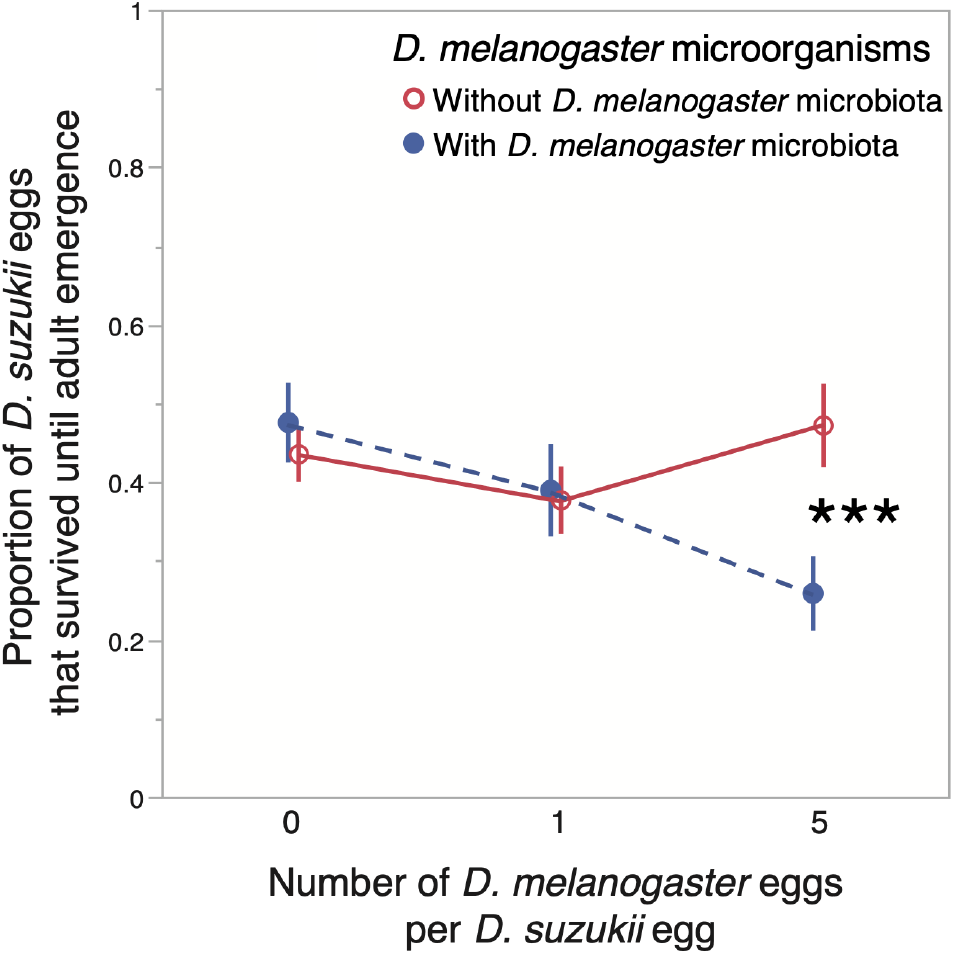
Effect of *D. melanogaster* larvae and their associated microbiota on the development of *D. suzukii* eggs until adult emergence. Eggs were individually deposited in grape berries where we mimicked natural oviposition by *Drosophila* females and field-like conditions. The greater ratio of *D. melanogaster* to *D. suzukii* egg follows relative infestation intensities observed in the field. The statistical interaction between number of *D. melanogaster* eggs and the presence or absence of their microbe was significant (F_2, 172_= 6.46; p= 0.002). Independent contrasts indicate a significant difference between the treatments with and without *D. melanogaster* microbes at high *D. melanogaster* density (F_1, 174_= 15.6; p= 0.0001). Overall REML model results: Number Dmel eggs per Ds egg; F_2, 165_= 4.83; p= 0.009; Dmel axenic or not; F_1, 162_= 0.41; p= 0.52; Number of Dmel eggs * axenic or not; F_2, 172_= 6.46; p= 0.002; Number of emerging Dmel adults; F_1, 174_= 7.74; p= 0.006. Symbols indicate means; error bars indicate standard-errors; *** for p< 0.001.

It Is remarkable that axenic *Dmel* larvae failed to reduce *Ds* larval development. This may provide an adaptive explanation for why *Ds* females did not respond to cues produced by axenic *Dmel. Ds* females should only avoid oviposition in environmental contexts that are detrimental to their offspring. The plastic decision by Ds to oviposit, or not, as a function of microbiological presence may enable the use of all suitable oviposition sites, with avoidance of sites only necessary when they are contaminated with costly competitors.

### Ecological significance and prospects for crop protection

Our study shows that commensal microbiota can mediate the competition between insect species with overlapping ecological niches. In our particular example, *Ds* females rely on combined cues from the competitor *Dmel* and its symbiont *L. brevis* to avoid oviposition sites that are likely to incur competition costs. It is well established that microorganisms impact the outcome of competitive interactions between hosts (29). Often, parasitic microorganisms shed by tolerant species have detrimental effects on less-tolerant competitors (e.g. 30); the spill-over hypothesis that facilitates the spread of some invasive species is based on this very mechanism (29). Symbiotic microorganisms can also elicit beneficial effects for heterospecific neighbors. For example, mycorrhizal fungi can mediate mutualism between plants species (31). In the present case, bacteria beneficial to the *Dmel* host are detrimental to *Ds* larvae, in complete opposition with how bacteria and yeast associated with *Ds* larvae facilitate fruit use by *Dmel* (24, 32). We hence document an original example of harmful interactions between competing insects mediated by their microbiota. Humans may exploit these interactions to protect the crops they grow.

Few species in the *Drosophila* genus oviposit in undamaged, ripening fruit. A phylogenetic perspective indicates that the ability to exploit ripening fruits is a derived character that evolved in *Ds* ancestors and presumably alleviates competition with other Drosophilids (33). *Dmel* arrived in Asia less than 60 000 years ago, long after the species origin of *Ds* (34). The larval niche of *Ds*, and possibly female oviposition preferences, hence probably evolved in response to other species of competitors. Several studies have reported that Ds larvae share their fruit with species such as as *Dmel, D. subobscura* and *Zaprionus indianus* in a variety of crop and wild plant species (24, 35, 36). In our experiments *Ds* did not avoid *D. simulans* cues. It is nonetheless plausible *Ds* females avoid cues produced by other Drosophilid species or populations, in particular those from the region it originates and possibly including other strains of *D. simulans*, and this avoidance may depend on the symbiotic status of those flies.

*Ds* is responsible for heavy crop losses throughout the globe due to the development of larvae in farmed fruit. It is tempting to exploit *Ds* oviposition avoidance to shelter fruit from *Ds* damage. Field-tests of repellents based on 1-octen-3-ol, a molecule produced by fungi that compete with *Drosophila* larvae, gave encouraging results (37, 38). In the present case, the microbiota associated with *Dmel* clearly cannot be sprayed directly in orchards because of the plastic avoidance loss exhibited by *Ds* females if they acquire those symbionts (Fig. 2a, 3c). A better solution may be to identify and use as a repellent the compounds produced by bacteria-inoculated *Dmel* (Fig. 3d). Future experiments would need to test whether *Ds* can become habituated to the aversive compound (39, 40) and whether management strategies such as refugia or alternating application need to be deployed. Characterizing *D. suzukii*’s chemosensory receptors and circuitry involved in the recognition of *Dmel* cues and its consequential behavioral response may enable the design of an optimized repellent.

## Materials and Methods

### General experimental design

The study is based on a simple assay where female *D. suzukii* (*Ds*) are given the choice to lay eggs on two substrates: either a blank control or a substrate that had previously been exposed to *D. melanogaster* (*Dmel*) adults (Fig. 1). By changing the nature of the *Ds* and *Dm* flies employed, we were able to reveal the factors that govern *Dmel*’s repellence and *Ds*’s corresponding avoidance.

In most cases, a single *Ds* female was placed in a 9cm diameter plastic cylindrical box for 24h. Boxes contained two 2*2cm*2cm plastic receptacles each half-filled with oviposition substrates, generally an agar-jellified strawberry puree or a piece of strawberry inserted in blank agar. These two substrates were prepared the day before, one of them was exposed to 3 adult *Dmel* flies overnight. Because these experiments were conducted over 5 years with variable objectives, some experimental details varied among assays. In all experiments, a variable fraction of assayed females (usually around 50%) did not oviposit during the 24h they spent with the tested substrates. These females were excluded from further analyses. Table 1 describes the experimental details, sample sizes and statistical analyses of each of the results reported in the article.

All flies were reared, and experiments conducted, in climatic chamber with a 13h:23°C/11h:19°C day/night cycle, an artificial dawn and dusk of 45min. Humidity was maintained constant at 75% relative humidity.

### Biological material

Most experiments were carried out with our standard *Ds* population that was founded by the authors in 2013 from a few dozen individuals that emerged from blackberries harvested in Gaujac, Southern France (44.0794, 4.5781); and the classical *Dmel* population Oregon R, founded in 1927 and shared among laboratories since then. These fly colonies were maintained in standard drosophila vials with banana artificial medium (see below) or 30 cm cubic cages when we needed larger numbers of flies.

Additional laboratory populations of *Ds* were as follows. The Japanese population was founded from individuals captured in Matsuyama, Japan (33.8389, 132.7917) in 2015 (courtesy A. Fraimout and V. Debat), the US population in Watsonville, California, USA (36.9144, -121.7577) in 2014 (individuals captured by S. F.), and the Chinese population in Shiping, China (23.7048, 102.5004) in 2015 (courtesy P. Girod and M. Kenis). The *D. simulans* population tested was founded from individuals captured in 2015 in Lyon, France (45.7835, 4.8791) (individuals captured by P. G.). All populations were initially composed of a few individuals and experienced repeated population bottlenecks during maintenance. They were thus largely inbred at the time of testing in 2017.

Wild *Ds* were captured during summer 2016 in two localities 10km apart near Montpellier, Southern France (43.6816, 3.8776), and tested about a week after capture, once they started laying eggs in the laboratory. Wild *Dmel* were also captured near Montpellier. For the experiment reported in Fig. 2d, *Dmel* flies were captured in several instances. Flies from a first group were reared in the laboratory and their offspring (*i*.*e*. F1) tested along with freshly-captured flies (*i*.*e*. F0). All wild flies were captured using custom-designed traps based on c.300 mL plastic cups, covered with cling-film, pierced on the sides for fly entry and containing an attractant (a mix water, vinegar, wine and sugar) separated from the flies by netting. The netting prevented fly drowning but allowed occasional access to the attractant as cups were readily shaken by wind or operators, which caused the netting to become soaked with the liquid bait. Traps were checked daily and usually contained various fly species, including *Dmel* and *Ds*.

### Recipes for rearing and oviposition media

Laboratory flies were reared on custom banana medium (1.2 L water, 280 g frozen organic banana, 74 g glucose, 74 g inactivated baker’s yeast, 12 g agar, 6 g paraben in 30 mL ethanol). The Chinese *Ds* population was reared in carrot medium (1.2 L water, 45 g carrot powder, 45 g glucose, 27 g inactivated baker’s yeast, 18 g corn meal, 13.5 g agar, 6 g paraben in 30 mL ethanol and 4 mL propionic acid).

In most cases, oviposition was assayed on strawberry puree (200 g frozen strawberry, 400 mL water, 6 g agar, 37 g glucose, 4 g paraben in 15 mL ethanol). In several instances (Table 1), we used jellified grape juice (100 mL commercial grape juice, 100 mL water, 12 g glucose, 2 g agar). Oviposition was also tested on pieces of strawberry inserted in jellified water (100 mL water, 1 g agar), they were first bleached (0.6% bleach during 5 min) to remove contaminants.

### Axenics, mono-associated flies and microbiological work

Axenic flies were produced following a protocol derived from (41). Briefly, *Drosophila* eggs were collected on grape-juice medium (see previous recipes section) before being bleached and rinsed twice (1.2% sodium hypochlorite). Eggs were then transferred to 50 mL centrifugation vials with 10 mL autoclaved banana medium (see recipes section) which lids were either incompletely screwed of harboured breathing membranes. All manipulations were conducted under a laminar flow hood. With care, it is possible to transfer freshly emerged adults to new vials aseptically and therefore maintain the population microbe-free for several generations. The axenic nature of the flies was regularly confirmed by the absence of cultivable microbes.

To produce mono-associated (i.e. gnotobiotic) adult flies, axenic flies were added to vials that had been surface-inoculated with suspensions (i.e. c. >10^5^ cells) of the relevant bacterium at least 4 days before experiment onset. The presence of inoculated microbes in adults was verified by culturing the bacteria retrieved from homogenised insects several days their nutritive medium was inoculated.

### Larval competition between D. suzukii and D. melanogaster in fruit

This assay aimed at testing whether the development of *Ds* larvae was affected by the presence of *Dmel* larvae and their associated microbiota. We took great care of reproducing field-like conditions (i.e. in-fruit interactions) as competition costs notoriously depend on ecological conditions (e.g. 42) and the effects of *Drosophila* bacterial symbionts on larval development change with medium richness (e.g. 43). A key parameter was to choose a fruit species in which both *Ds* and *Dmel* had been reported to develop simultaneously in the field, and we elected grape (24). Given the large effect of grape variety on Ds oviposition (44), we first confirmed that *Ds* would oviposit on the batch of grapes we used (fruit of an unknown cultivar bought in April 2018 in a food retail store) and that this behaviour was reduced by exposure to *Dmel* (data not shown). In order to mimic realistic competition conditions we manually pierced the skin of grape berries with fine needles, making a hole close in size as those *Ds* females do with their ovipositor (33). Each hole was first inoculated with a wild strain of the yeast *Hanseniaspora meyeri* isolated from wild *Ds* adults and received a single *Ds* egg (Fig. S3a). There were 6 holes per berry. Each of these holes also received 0, 1 or 5 *Dmel* eggs. The larger ratio of *Dmel* to *Ds* eggs was chosen as it reflects relative infestation intensity observed in grapes collected in the field (e.g. 24). In half the cases, deposited *Dmel* eggs had been made axenic by bleaching (see previous section on the production of axenic flies). Note that an important design choice was to either compare the effect of axenic and conventional *Dmel* larvae, or axenic *Dmel* and axenic *Dmel* artificially inoculated with microbes harvested from conventional flies. We rejected the second option because it would have been impossible to been certain that eggs artificially associated to microorganisms cocktails bore all relevant strains. By contrast, the differential mortality of bleached (i.e. axenic) and non-bleached (i.e. conventional) eggs could be controlled for statistically (see statistical analysis section below; Fig. S3b) In the treatments without *Dmel* eggs but with its microbiota, pierced berries were exposed to 10 *Dmel* males for 24h prior to *Ds* egg deposition. All grape berries were incubated in individual plastic vials until adult flies emerged. This assay comprised 25-30 individual berries per treatment (50 replicates for the control treatment with *Ds* eggs and no *Dmel* microbiota) spread over 8 temporal blocks.

### Statistical analyses

In all reported experiments except the one on larval competition (Fig. 4), *Ds* females deposited their eggs on either treated or untreated oviposition substrates. Egg counts on each type of medium were therefore not independent because were produced by the same females. Additionally, total number of eggs varied among females and experiments and largely followed a Poisson distribution, which prevented the use of traditional linear models that assume normal distributions of the residuals. We therefore used a simple, robust statistical approach to analysing the proportion of eggs deposited on treated and untreated site: a non-parametric, one-tailed Wilcoxon signed rank test that took into account data pairing, was compatible with the data distribution, and has often been used in comparable studies (e.g.13). We noticed that paired t-tests, which assume data follow a normal distribution, provided similar results. The aim of our experiments was to investigate female behaviour determinants rather than infestation intensities, so the units of replication were the females and their individual preferences towards different types of substrates. For this reason, the statistical methods we employed were not affected by variation in the fecundity of individual females, and the most fertile females could not skew the results towards their specific preferences. With this in mind, it appeared preferable to include all females that oviposited, even if those that deposited only a single egg. Because of the plasticity of the avoidance behaviour, all experiments included a positive control - usually the response of standard *Ds* to laboratory *Dmel* flies. This ensured that lack of avoidance in an experiment was not due to unidentified factors or inappropriate conditions. Note that several of our most important results were repeatedly observed in distinct experiments. Compare, for example, loss of avoidance in Fig. 2a and Fig. 3c, effect of axenic *Dmel* in Fig. 3a and Fig. 3b, restauration of *Dmel* repellency by *Lactobacillus brevis* inoculation in Fig. 3b and Fig. 3c.

Results from the larval competition assay were analysed using a linear mixed-model with the REML method. Numbers of *Ds* adult that emerged from each fruit were Log(x+1)-transformed and complied with tests assumptions. This model contained discreet, fixed terms describing the number of *Dmel* eggs deposited, whether *Dmel* microbiota was present, and their interaction. It was also very important that the model included the (log-transformed) number of *Dmel* adults that emerged from the fruit as a fixed, continuous factor. Indeed, this term was necessary to control for the additional mortality of *Dmel* larvae caused by bleaching eggs in the axenic treatment (Fig. S3b). The presence of this term in the analysis ensures the significant effect of axeny was not an artefact due to reduced *Dmel* larvae numbers. The model also included a block term (treated as random). Differences among treatments were tested with independent contrasts and pairwise student’s tests.

All analyses were carried out with the software JMP 14.0 (SAS Institute Inc. 2018). Throughout the manuscript, stars in figures indicate significance of one-tailed statistical tests: * p<0.05; ** p<0.01; ***p<0.001; n.s. p>0.05.

All data is available on the Zenodo platform under the reference: 10.5281/zenodo.3970737.

## Acknowledgements

We are grateful to Marc Kenis and Pierre Girod for providing the Chinese *D. suzukii* population; Antoine Fraimout and Vincent Debat for the Japanese *D. suzukii* population; Marie-Pierre Chapuis and Laure Benoit for help with microbial work.

## References

1. Fraimout A, Debat V, Fellous S, Hufbauer R, Foucaud J, Pudlo P, et al. Deciphering the routes of invasion of Drosophila suzukii by means of ABC random forest. Mol Biol Evol. 2017.

2. De Ros G, Conci S, Pantezzi T, Savini G. The economic impact of invasive pest Drosophila suzukii on berry production in the Province of Trento, Italy. Journal of Berry Research. 2015;5(2):89–96.

3. Yeh DA, Drummond FA, Gómez MI, Fan X. The Economic Impacts and Management of Spotted Wing Drosophila (Drosophila Suzukii): The Case of Wild Blueberries in Maine. Journal of Economic Entomology. 2020;113(3):1262–9.

4. Mazzi D, Bravin E, Meraner M, Finger R, Kuske S. Economic Impact of the Introduction and Establishment of Drosophila suzukii on Sweet Cherry Production in Switzerland. Insects. 2017;8(1).

5. Dancau T, Stemberger TL, Clarke P, Gillespie DR. Can competition be superior to parasitism for biological control? The case of spotted wing Drosophila (Drosophila suzukii), Drosophila melanogaster and Pachycrepoideus vindemmiae. Biocontrol Science and Technology. 201727(1):3–16.

6. Shaw B, Brain P, Wijnen H, Fountain MT. Reducing Drosophila suzukii emergence through inter-species competition. Pest management science. 2017.

7. Elsensohn JE, Aly MF, Schal C, Burrack HJ. Social signals mediate oviposition site selection in Drosophila suzukii. Scientific Reports. 2021;11(1):1–10.

8. Tait G, Park K, Nieri R, Crava MC, Mermer S, Clappa E, et al. Reproductive Site Selection: Evidence of an Oviposition Cue in a Highly Adaptive Dipteran, Drosophila suzukii (Diptera: Drosophilidae). Environmental Entomology. 2020;49(2):355–63.

9. Asplen MK, Anfora G, Biondi A, Choi D-S, Chu D, Daane KM, et al. Invasion biology of spotted wing Drosophila (Drosophila suzukii): a global perspective and future priorities. J Pest Sci. 2015;88(3):469–94.

10. Silva MA, Bezerra-Silva GCD, Mastrangelo T. The host marking pheromone application on the management of fruit flies-a review. Brazilian Archives of Biology and Technology. 2012;55(6):835–42.

11. Capy P, Gibert P. Drosophila Melanogaster, Drosophila Simulans: so Similar yet so Different. Genetica. 2004;120(1):5–15.

12. Stensmyr Marcus C, Dweck Hany KM, Farhan A, Ibba I, Strutz A, Mukunda L, et al. A Conserved Dedicated Olfactory Circuit for Detecting Harmful Microbes in Drosophila. Cell. 2012;151(6):1345–57.

13. Wong AC-N, Wang Q-P, Morimoto J, Senior AM, Lihoreau M, Neely GG, et al. Gut microbiota modifies olfactory-guided microbial preferences and foraging decisions in Drosophila. Curr Biol. 2017;27(15):2397-404. e4.

14. Qiao H, Keesey IW, Hansson BS, Knaden M. Gut microbiota affects development and olfactory behavior in Drosophila melanogaster. Journal of Experimental Biology. 2019;222(5).

15. Chandler JA, James PM, Jospin G, Lang JM. The bacterial communities of Drosophila suzukii collected from undamaged cherries. PeerJ. 20142:e474.

16. Staubach F, Baines JF, Künzel S, Bik EM, Petrov DA. Host species and environmental effects on bacterial communities associated with Drosophila in the laboratory and in the natural environment. PloS one. 2013;8(8):e70749.

17. Wang Y, Kapun M, Waidele L, Kuenzel S, Bergland AO, Staubach F. Common structuring principles of the Drosophila melanogaster microbiome on a continental scale and between host and substrate. Environmental Microbiology Reports. 2020;12(2):220–8.

18. Chandler JA, Morgan Lang J, Bhatnagar S, Eisen JA, Kopp A. Bacterial Communities of Diverse Drosophila Species: Ecological Context of a Host–Microbe Model System. PLoS Genet. 2011;7(9):e1002272.

19. Ryu J-H, Kim S-H, Lee H-Y, Bai JY, Nam Y-D, Bae J-W, et al. Innate Immune Homeostasis by the Homeobox Gene Caudal and Commensal-Gut Mutualism in Drosophila. Science. 2008;319(5864):777.

20. Shin SC, Kim S-H, You H, Kim B, Kim AC, Lee K-A, et al. Drosophila Microbiome Modulates Host Developmental and Metabolic Homeostasis via Insulin Signaling. Science. 2011;334(6056):670.

21. Tracy C, Krämer H. Escherichia coli Infection of Drosophila. Bio-protocol. 2017;7(9).

22. Sato A, Tanaka KM, Yew JY, Takahashi A. Drosophila suzukii avoidance of microbes in oviposition choice. Royal Society Open Science. 2021;8(1):201601.

23. Li H, Ren L, Xie M, Gao Y, He M, Hassan B, et al. Egg-Surface Bacteria Are Indirectly Associated with Oviposition Aversion in Bactrocera dorsalis. Curr Biol. 2020.

24. Rombaut A, Guilhot R, Xuéreb A, Benoit L, Chapuis M-P, Gibert P, et al. Invasive Drosophila suzukii facilitates Drosophila melanogaster infestation and sour rot outbreaks in the vineyards. Royal Society Open Science. 2017.

25. Bing X, Gerlach J, Loeb G, Buchon N. Nutrient-Dependent Impact of Microbes on Drosophila suzukii Development. mBio. 2018;9(2).

26. Guilhot R, Rombaut A, Xuéreb A, Howell K, Fellous S. Environmental specificity in Drosophila-bacteria symbiosis affects host developmental plasticity. Evol Ecol. 2020;34:693– 712

27. Khodaei L, Long TAF. Kin Recognition and Egg Cannibalism by Drosophila melanogaster Larvae. Journal of Insect Behavior. 2020;33(1):20–9.

28. Vijendravarma RK, Narasimha S, Kawecki TJ. Predatory cannibalism in Drosophila melanogaster larvae. Nature Communications. 2013;4(1):1789.

29. Amsellem L, Brouat C, Duron O, Porter SS, Vilcinskas A, Facon B. Importance of microorganisms to macroorganisms invasions: is the essential invisible to the eye?(The Little Prince, A. de Saint-Exupéry, 1943). Advances in Ecological Research. 57: Elsevier; 2017. p. 99–146.

30. Tompkins D, White AR, Boots M. Ecological replacement of native red squirrels by invasive greys driven by disease. Ecol Lett. 2003;6(3):189–96.

31. Selosse M-A, Richard F, He X, Simard SW. Mycorrhizal networks: des liaisons dangereuses? Trends Ecol Evol. 2006;21(11):621–8.

32. Solomon GM, Dodangoda H, McCarthy-Walker T, Ntim-Gyakari R, Newell PD. The microbiota of Drosophila suzukii influences the larval development of Drosophila melanogaster. PeerJ. 2019;7:e8097.

33. Atallah J, Teixeira L, Salazar R, Zaragoza G, Kopp A. The making of a pest: the evolution of a fruit-penetrating ovipositor in Drosophila suzukii and related species. Proceedings of the Royal Society of London B: Biological Sciences. 2014;281(1781):20132840.

34. Stephan W, Li H. The recent demographic and adaptive history of Drosophila melanogaster. Heredity. 2007;98(2):65–8.

35. Bernardi D, Andreazza F, Botton M, Baronio C, Nava D. Susceptibility and Interactions of Drosophila suzukii and Zaprionus indianus (Diptera: Drosophilidae) in Damaging Strawberry. Neotropical Entomology. 2016:1–7.

36. Poyet M, Eslin P, Héraude M, Le Roux V, Prévost G, Gibert P, et al. Invasive host for invasive pest: when the Asiatic cherry fly (Drosophila suzukii) meets the American black cherry (Prunus serotina) in Europe. Agricultural and Forest Entomology. 2014:n/a-n/a.

37. Wallingford AK, Cha DH, Linn JCE, Wolfin MS, Loeb GM. Robust Manipulations of Pest Insect Behavior Using Repellents and Practical Application for Integrated Pest Management. Environmental Entomology. 2017;46(5):1041–50.

38. Wallingford AK, Connelly HL, Dore Brind’Amour G, Boucher MT, Mafra-Neto A, Loeb GM. Field Evaluation of an Oviposition Deterrent for Management of Spotted-Wing Drosophila, Drosophila suzukii, and Potential Nontarget Effects. Journal of Economic Entomology. 2016.

39. Kahsai L, Zars T. Learning and memory in Drosophila: behavior, genetics, and neural systems. International review of neurobiology. 99: Elsevier; 2011. p. 139–67.

40. Little CM, Chapman TW, Hillier NK. Considerations for Insect Learning in Integrated Pest Management. Journal of Insect Science. 2019;19(4).

41. Koyle ML, Veloz M, Judd AM, Wong AC-N, Newell PD, Douglas AE, et al. Rearing the fruit fly Drosophila melanogaster under axenic and gnotobiotic conditions. JoVE (Journal of Visualized Experiments). 2016(113):e54219.

42. Kraaijeveld AR, Godfray HC. Trade-off between parasitoid resistance and larval competitive ability in Drosophila melanogaster. Nature. 1997;389(6648):278–80.

43. Storelli G, Defaye A, Erkosar B, Hols P, Royet J, Leulier F. Lactobacillus plantarum Promotes Drosophila Systemic Growth by Modulating Hormonal Signals through TOR-Dependent Nutrient Sensing. Cell Metabolism. 2011;14(3):403–14.

44. Mazzetto F, Lessio F, Giacosa S, Rolle L, Alma A. Relationships between Drosophila suzukii and grapevine in North-western Italy: seasonal presence and cultivar susceptibility. Bulletin of Insectology. 2020;73(1):29–38.

