## Supplementary material for "Microbiota-Mediated Competition Between *Drosophila* Species"

### S1. *D. suzukii* females were neither attracted nor repelled by sites with eggs from conspecifics

Shaw *et al.* (1) reported *D. suzukii* (*Ds*) females were neither attracted nor repelled by sites with conspecific eggs. We repeatedly observed the same pattern, as in the experiment described below. Ovipositional preference of *Ds* females was tested with 9cm petri-dishes with artificial medium that already contained, or not, naturally-deposited conspecific eggs (between 19 and 147). The experiment was conducted in 30 cm diameter cylinder cages with 10 females over two successive days keeping the same females. Effect of conspecific eggs presence was not significant (Wilcoxon signed rank tests) ; it was not influence by the prior number of eggs present in the medium (RMEL mixed-effect model on proportion of eggs deposited on each type of substrate: $F_{1,13}$= 1.37, p= 0.26).

Recent literature has however revealed *DS* females can deposit marking cues during oviposition (2, 3). These cues sometimes attract oviposition by conspecifics (3) but they also repel DS (2). (2) argue these contradictory observations would be explained by the context-dependency of DS oviposition preferences.

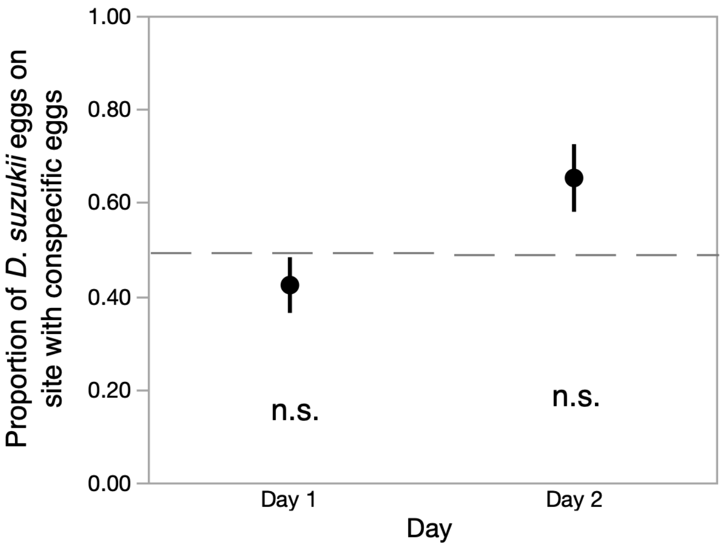


Fig. S1: presence of *D. suzukii* eggs did not elicit *D. suzukii* oviposition avoidance. Wilcoxon signed rank tests were not significant on either day. Symbols indicate means and error-bars standard errors.

### S2. Short-range repellence by *D. melanogaster*

In order to test whether *D. melanogaster* (*Dmel*) repellency could proceed at a distance, for example through odorant volatiles, we designed an assay based on the one of (4). We used the same 2-cm cubic receptacles as for most oviposition assays, which we exposed to *Dmel* for 24h (control media were not exposed to *Dmel*). These receptacles were placed with the aperture on the side on 12cm square petri dishes containing jellified grape juice within a 20 cm cubic netted cage. The assay was then conducted as usual with single *Ds* females from our standard population. After 24h, we counted the number of eggs on the medium in the small receptacle that contained exposed (or unexposed) strawberry puree as well as on the surrounding medium.

As expected, exposed media received less *Ds* eggs than unexposed ones. However, the effect did not extend to the surrounding substrates (Fig. S2). This shows that repellency does not rely on long-range odors and suggests gustation and direct contact may be required by *Ds* females to assess *Dmel* cues.

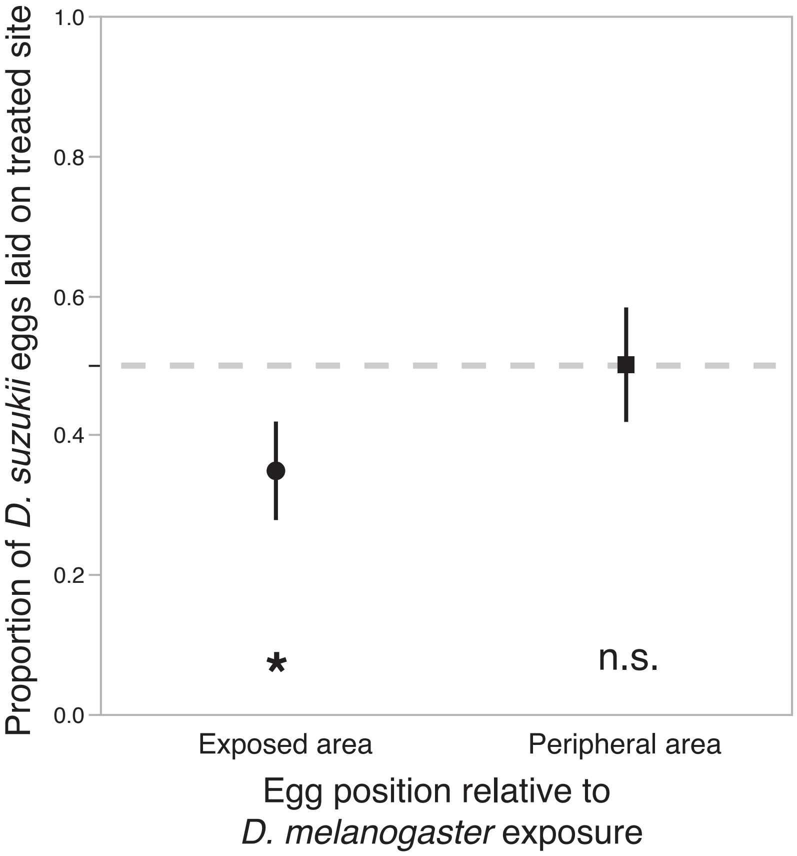


Fig. S2: fresh medium (right) placed close to medium previously exposed to *D. melanogaster* (left) is not avoided by egg-laying *D. suzukii* females. Symbols indicate means and error-bars standard errors. Statistical tests produced by Wilcoxon signed rank tests; * for p< 0.05.

Table S2: methodological and statistical details.

| **Experiment** question and figure presenting the results | **Method** | **Number of *Ds* females in assay; type of container** | ***Ds* particulars** | ***Dmel* particulars** | **Oviposition substrate** | **Replication and raw statistical results** Reported number of replicates excludes the frequent cases where no *Ds* eggs were deposited during the experiment | **Additional comments** |
|---|---|---|---|---|---|---|---|
| - Is *Dmel* repellence exerted at a distance?<br>- Fig. S2 | We tested whether medium surrounding medium exposed to *Dmel* received less *Ds* eggs than medium surrounding pristine medium. | - 1 female per assay<br>- 20cm cubic netting cages | Standard laboratory population | Standard Oregon R laboratory population | - strawberry puree<br>- 2*2cm cubic receptacles and 12cm square petri dishes | **Wilcoxon signed rank tests, one-tailed:**<br>Exposed area: n= 29, S= 101; p= 0.013<br>Peripheral area: n= 29, S= -14; p= 0.61 | Details of the protocol are described in the supplementary materials |

### S3. Protocol for within-fruit competition between larvae

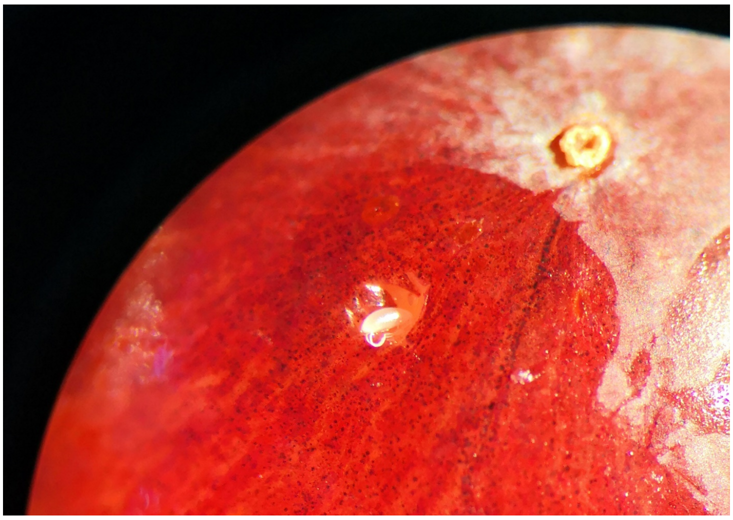

Fig. S3a: A single *D. suzukii* egg manually deposited in an artificial oviposition hole. In the experiment reported here, fruit bore 6 such eggs. In some treatments, 1 or 5 *D. melanogaster* eggs were also deposited in the hole. In half of these treatments, *D. melanogaster* eggs had been made axenic using bleach baths.

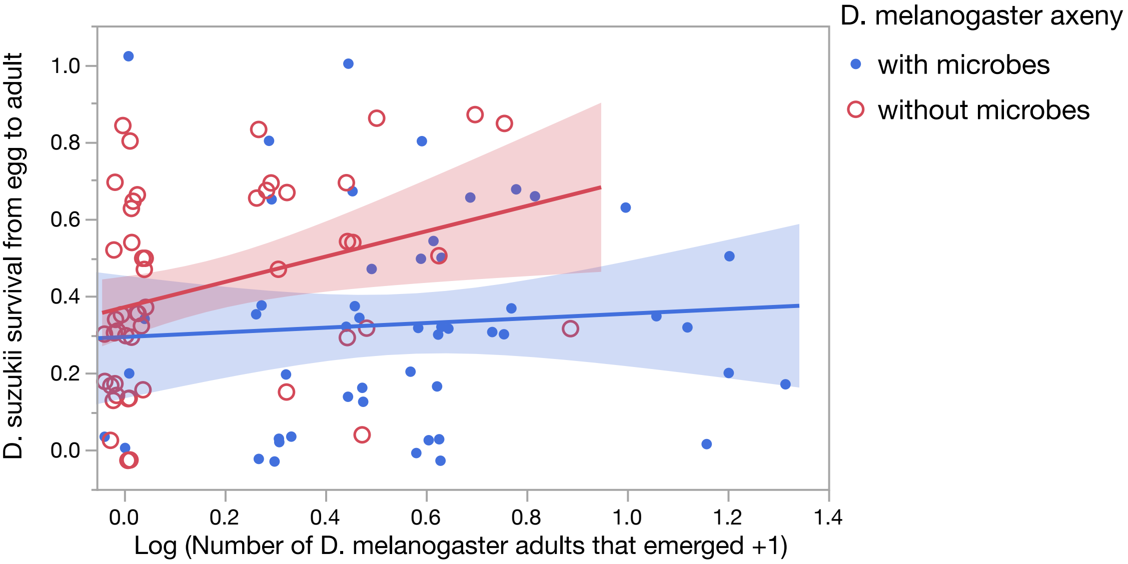


Fig. S3b: relationship between the number of live *D. melanogaster* present in the fruit and *D. suzukii* survival from larval to adult stage. Numbers of emerging *Dmel* were included in statistical analyses of *D. suzukii* development (Fig. 5) in order to control for additional *D. melanogaster* mortality induced by egg bleaching in the axenic treatment. Note that when *D. melanogaster* adults emerged in similar numbers in the axenic and conventional treatments (i.e. blue and red dots in the central part of the X axis), *D. suzukii* survival was superior in absence of *D. melanogaster* microbes. In this figure, different regressions were fitted for axenic and conventional treatments even though the interaction between *D. melanogaster* axeny treatment and the X axis was not significant ($F_{1,172}$= 0.036, P= 0.85). Points were slightly jittered for readability.

**References for Supplementary Materials**

1. Shaw B, Brain P, Wijnen H, Fountain MT. Reducing *Drosophila suzukii* emergence through inter-species competition. Pest management science. 2017.

2. Elsensohn JE, Aly MF, Schal C, Burrack HJ. Social signals mediate oviposition site selection in *Drosophila suzukii*. Scientific Reports. 2021;11(1):1-10.

3. Tait G, Park K, Nieri R, Crava MC, Mermer S, Clappa E, et al. Reproductive Site Selection: Evidence of an Oviposition Cue in a Highly Adaptive Dipteran, *Drosophila suzukii* (Diptera: Drosophilidae). Environmental Entomology. 2020;49(2):355-63.

4. Karageorgi M, Bräcker LB, Lebreton S, Minervino C, Cavey M, Siju KP, et al. Evolution of Multiple Sensory Systems Drives Novel Egg-Laying Behavior in the Fruit Pest *Drosophila suzukii*. Curr Biol. 2017;27(6):847-53.
